# Shrimp parvovirus circular DNA fragments arise from both endogenous viral elements (EVE) and the infecting virus

**DOI:** 10.1101/2021.06.08.447433

**Authors:** Suparat Taengchaiyaphum, Phasini Buathongkam, Suchitraporn Sukthaworn, Prapatsorn Wongkhaluang, Kallaya Sritunyalucksana, Timothy William Flegel

## Abstract

Some insects use endogenous reverse transcriptase (RT) to make variable viral copy DNA (vcDNA) fragments from viral RNA in linear (lvcDNA) and circular (cvcDNA) forms. The latter form is easy to extract selectively. The vcDNA produces small interfering RNA (siRNA) variants that inhibit viral replication via the RNA interference (RNAi) pathway. The vcDNA is also autonomously inserted into the host genome as endogenous viral elements (EVE) that can also result in RNAi. We hypothesized that similar mechanisms occurred in shrimp. We used the insect methods to extract circular viral copy DNA (cvcDNA) from the giant tiger shrimp (*Penaeus monodon*) infected with a virus originally named infectious hypodermal and hematopoietic necrosis virus (IHHNV). Simultaneous injection of the extracted cvcDNA plus IHHNV into whiteleg shrimp (*Penaeus vannamei*) resulted in a significant reduction in IHHNV replication when compared to shrimp injected with IHHNV only. Next generation sequencing (NGS) revealed that the extract contained a mixture of two general IHHNV-cvcDNA types. One showed 98 to 99% sequence identity to GenBank record AF218266 from an extant type of infectious IHHNV. The other type showed 98% sequence identity to GenBank record DQ228358, an EVE formerly called non-infectious IHHNV. The startling discovery that EVE could also give rise to cvcDNA revealed that cvcDNA provided an easy means to identify and characterize EVE in shrimp and perhaps other organisms. These studies open the way for identification, characterization and use of protective cvcDNA as a potential shrimp vaccine and as a tool to identify, characterize and select naturally protective EVE to improve shrimp tolerance to homologous viruses in breeding programs.

## 1. INTRODUCTION

In 2009 (1), it was hypothesized that endogenous viral elements (EVE) with high sequence identity to extant viruses in shrimp and insects arise via host recognition of viral messenger RNA followed by formation of variable cDNA fragments (here called viral copy DNA or vcDNA) from it by host reverse transcriptase (RT). Integration of those vcDNA fragments into the host genome is via host integrase (IN). The EVE give rise to negative sense RNA that result in degradation of viral RNA by the RNA interference (RNAi) pathway. It was proposed that this is the underlying natural mechanism that leads to balanced persistent infections in which one or more viruses are tolerated by shrimp and insects, sometimes for a lifetime, without signs of disease. This phenomenon of tolerance to persistent viral infections had been called viral accommodation (2–4) but the underlying mechanisms involving EVE were not hypothesized until 2009 (1, 5). Viral accommodation via EVE constitutes a process of autonomous genetic modification (AGMo) that gives rise to natural transgenic organisms (NTO), and accommodation is heritable if the EVE occur in germ cells. Some predictions of the hypothesis have been supported by research on insects since 2013 (6–11) and proof of a protective EVE against a virus in mosquitoes was published in 2020 (12). An updated summary diagram of the currently hypothesized mechanisms related to viral accommodation is shown in **Figure 1**.

**Figure 1.**
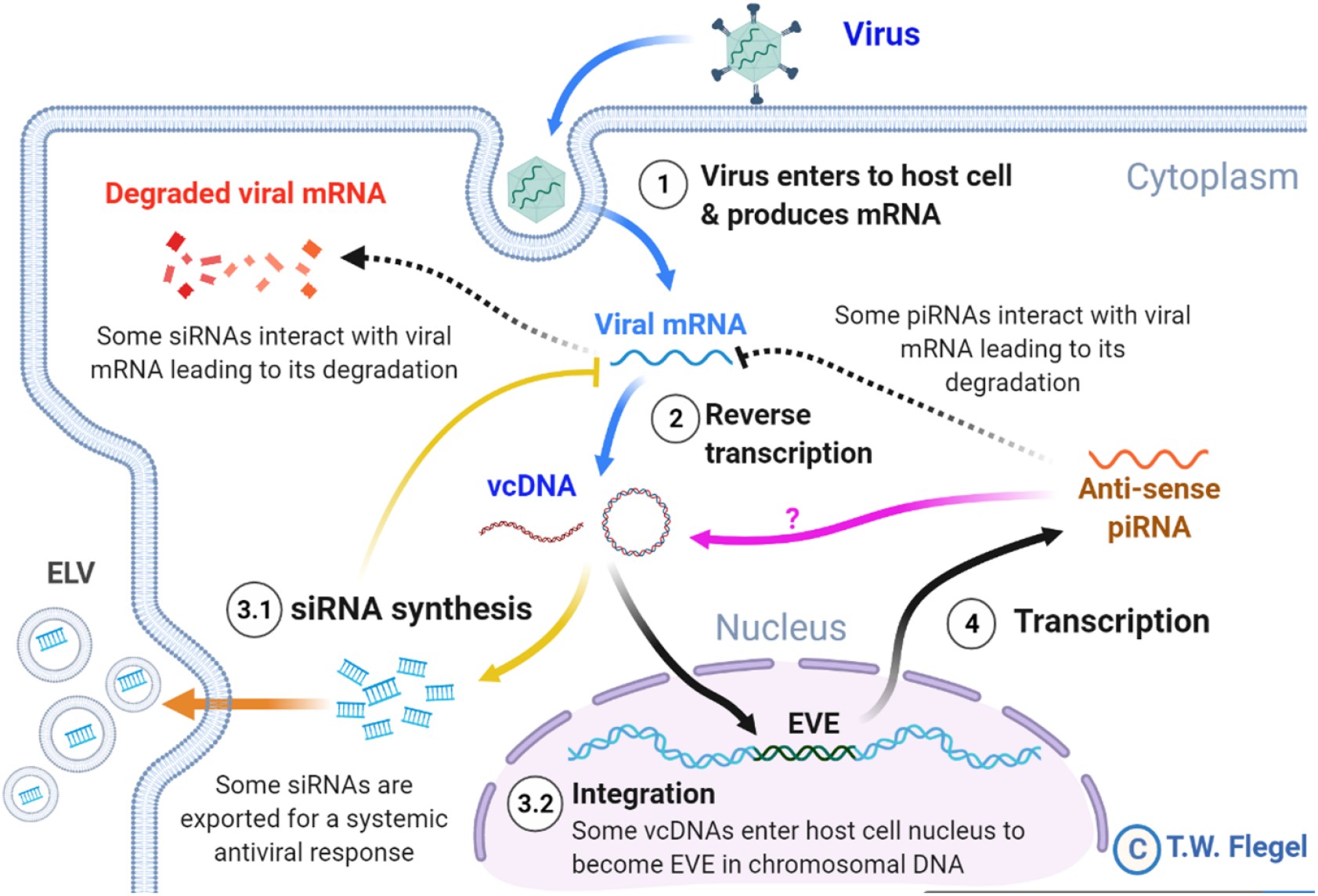
A simplified diagram of the mechanisms involved in viral accommodation as updated from Flegel (2020). The update includes additional pathways (indicated by yellow and orange arrows) that were not foreseen in the 2009 viral accommodation hypothesis (blue and black arrows). Specifically, vcDNA was not predicted to occur also in a circular form (cvcDNA). In addition, no immediate production of siRNA leading to an RNAi response was predicted. Nor was the occurrence of exosome like vesicles (ELV) for systemic dispersal of the RNAi response predicted. Nor was the discovery (this paper) that EVE could produce viral circular DNA (purple arrow). All these features are hypothesized to occur in shrimp. Abbreviations used: siRNA for small interfering RNA and piRNA for PIWI-interacting RNA(s).

Not predicted by the viral accommodation hypothesis of 2009 was the discovery that vcDNA produced by the action of host RT upon viral infection occurs in both linear (lvcDNA) and circular (cvcDNA) forms that, in turn, immediately produce small interfering RNA (siRNA) transcripts that result in an immediate and specific cellular and systemic RNAi response to invading viruses (7, 8, 10). Although all these new discoveries were made using RNA virus models, we considered it possible that they might also occur in shrimp since they too have been reported to have EVE homologous to extant DNA viruses (13–15). We were particularly interested in cvcDNA and the possibility that shrimp would produce protective cvcDNA in a manner similar to that reported for insects (10). We hypothesized that use of the techniques devised for extraction of cvcDNA from insects would be successful when used with the giant tiger shrimp (*Penaeus monodon*) infected with *Penstylhamaparvovirus* 1 from the family *Parvoviridae* and sub-family *Hamaparvovirinae* (16). This single-stranded DNA virus was previously called infectious hypodermal and hematopoietic necrosis virus (IHHNV) and we will use that acronym here to maintain easy links to previous literature. We also hypothesized that the extracted cvcDNA would significantly reduce IHHNV replication in the whiteleg shrimp *Penaeus vannamei* challenged with IHHNV.

## 2. MATERIALS AND METHODS

### 2.1. PCR methods and primers used in this study

The PCR primers used in this study are shown in **Table 1.** To determine the presence of infectious IHHNV and to test its replication level in challenged shrimp, a long-amp IHHNV detection method was used to detect a 3665 base-region of IHHNV (approximately 92% of the whole genome and excluding its hairpin ends). A Long-Amp^™^ *Taq* PCR mix (New England Biolab, USA) was used. The PCR reaction consisted of Long-Amp *Taq* PCR reaction mix, 0.4 μM of forward and reverse primers (98F/3762R), 1U Long-Amp^™^ *Taq* polymerase, and either 100 ng DNA before digestion or 2 ng DNA post-enzyme digestion. The PCR cycle was started with initial denaturation at 94°C for 30 s then followed by 35 cycles of 94°C for 20 s, 55°C for 30 s, 72°C for 2.5 min and final extension at 72°C for 10 min.

**Table 1.**
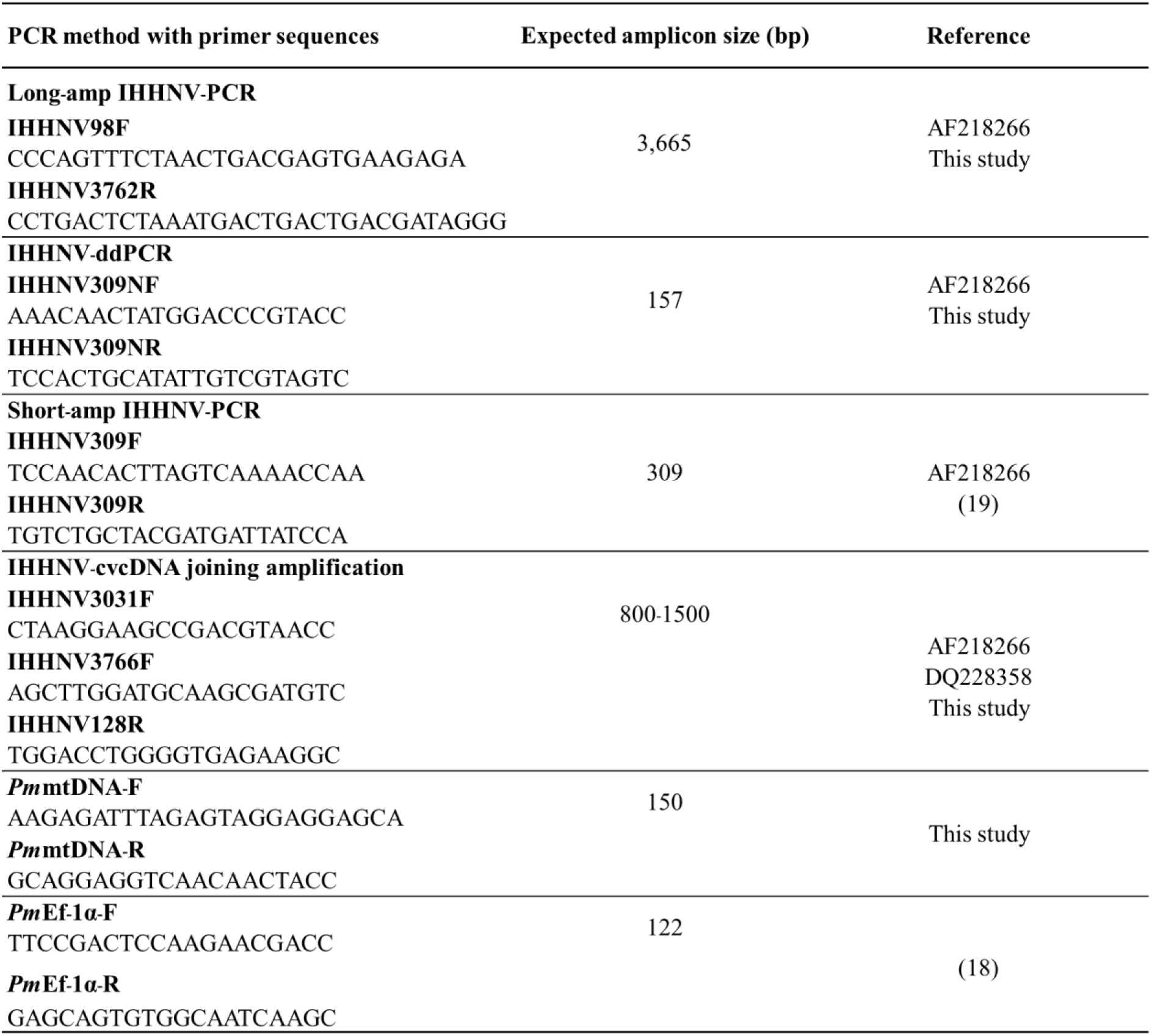
List of the primers used in this study.

Quantitative PCR by droplet digital PCR (IHHNV-ddPCR) was used to check the number of viral copies in the crude IHHNV stock and the number of IHHNV-cvcDNA in the circular DNA preparation. The ddPCR reaction was prepared by using EvaGreen^™^ ddPCR supermix (Bio-Rad, USA) which consisted of 1X ddPCR mix, 0.2 μM of forward and reverse primers (309NF/309NR), and either 1 μl of diluted crude viral stock (at 10^-7^ dilution) or 1 ng of the circular DNA preparation as the template. The ddPCR amplification cycle was set according to the manufacturer’s protocol by adjusting the annealing temperature to 56°C. After the complete PCR cycles, the reactions were analyzed by fluorescent signal using a ddPCR plate reader. The absolute amount of target DNA copy per reaction was calculated based on Pearson’s correlation method using QuantaSoft^™^ ddPCR analysis software (Bio-Rad, USA). PCR reactions for each individual sample were performed in duplicate.

The short-amp IHHNV-PCR method (17) was used to check for infectious IHHNV sequences in DNA extracts and in infected shrimp. As an internal control gene for linear, chromosomal DNA, primers specific to shrimp elongation factor 1 alpha (EF-1α) gene were used to give an amplicon of 122 bp (18). PCR amplicons were analyzed by 1.5% agarose gel electrophoresis followed by visualization of ethidium bromide staining by UV light.

### 2.2 Preparation of crude IHHNV stock inoculum

Frozen black tiger shrimp (*Penaeus monodon*) were checked for IHHNV infection using the long amp- IHHNV detection method. The pleopods from 5 IHHNV-positive shrimp were collected, pooled, homogenized and dissolved in cold 1X PBS pH 7.4. The tissue homogenate was centrifuged at 8,000 rpm to remove cell debris before it was subjected to filtration through a 0.2 μm membrane filter. The filtrate was collected and aliquoted into small tubes and referred to as “crude IHHNV stock”. A similar protocol was applied to 5 IHHNV-negative shrimp (*P. monodon*) samples from the same batch of frozen shrimp. The crude IHHNV stock was subsequently used in the challenge tests with *P. vannamei* where IHHNV infection and replication was confirmed by PCR. The crude IHHNV stock was stored at - 80°C for further experiments. To check the virus titer, crude IHHNV stock was serially diluted and subjected to quantification by the IHHNV-ddPCR.

The sequence of the long-amp IHHNV amplicon in the DNA extract from the crude IHHNV stock was determined and compared with previously reported IHHNV sequences in the Genbank database using the UPGMA method and MEGA X software (https://www.megasoftware.net/). Results revealed that the crude IHHNV stock contained a type of IHHNV closely related to isolates previously reported from Thailand (AY362547 and AY102034) and Taiwan (AY355307) (see **Figure S1)**.

**Figure S1.**
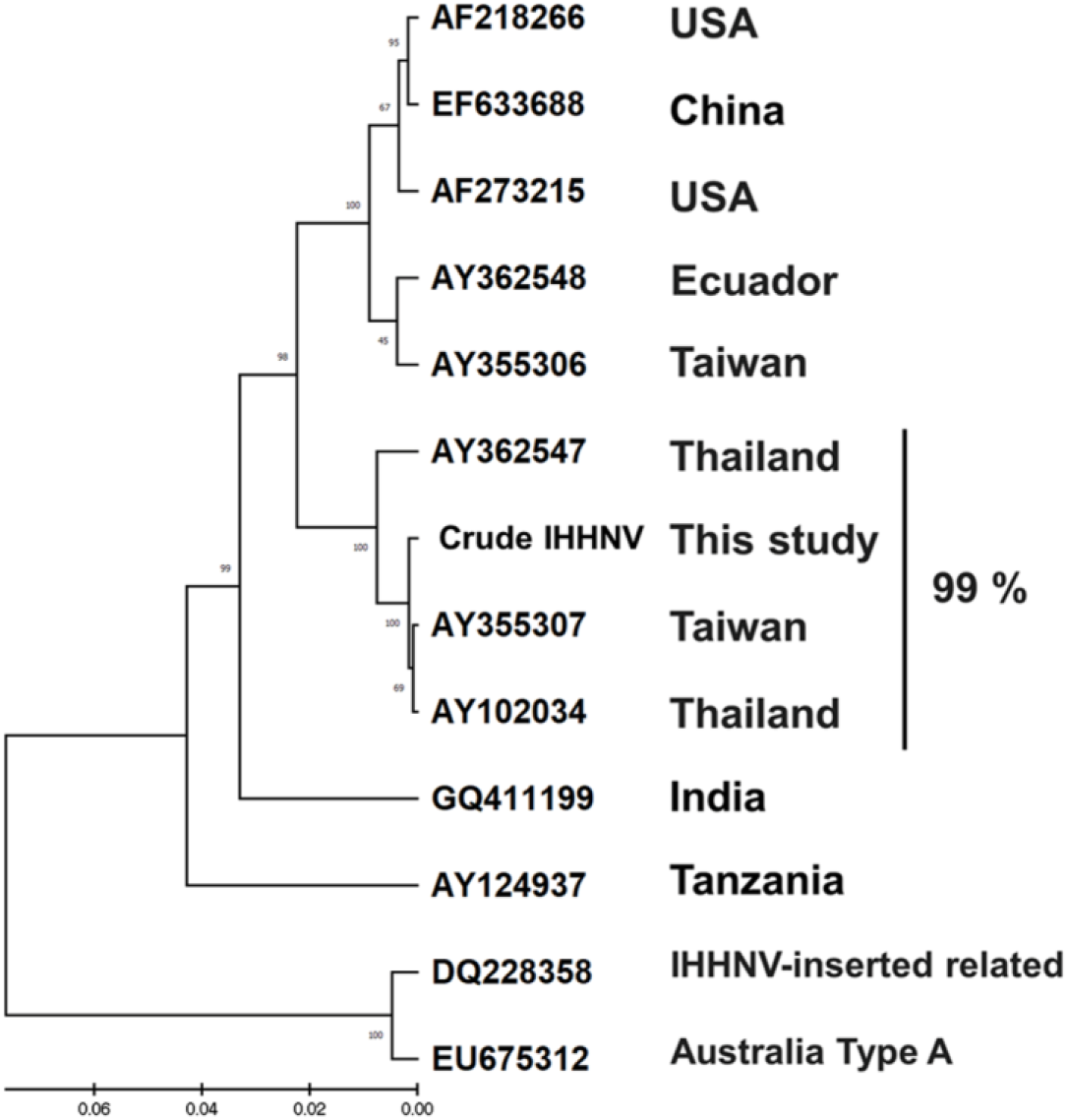
UPGMA cluster tree constructed for comparison of 2814 bases of our nucleic acid sequence (GQ475529) with matching regions of IHHNV sequences at GenBank. The scale bar below the tree indicates the relative difference in percent identity. The vertical bars on the right of the tree indicate the percentage or percentage range of differences in identity among the isolates covered by the bars. Since the sequences of AY355308, and AY355306, <u>AY355308</u>, <u>AY355306</u> were identical in the region compared, only the sequence of <u>AY355306</u> was used in constructing the tree.

After receiving the NGS sequencing results for the cvcDNA preparations from IHHNV-infected shrimp, the DNA extracts and the cvcDNA preparations from the IHHNV-negative and IHHNV-positive *P. monodon* were tested by PCR (19) for the presence of an IHHNV-EVE corresponding to the GenBank reference sequence DQ228358.

### 2.3 Extraction of circular DNA from IHHNV-infected shrimp

Total DNA extract from pleopods of the IHHNV-infected *P. monodon* samples was subjected to circular DNA isolation as previously described (10). Briefly, total shrimp DNA was prepared using a DNA extraction kit (Qiagen, USA) and DNA concentration was determined by NanoDrop spectrophotometer (Thermo Scientific, USA). Extraction of circular DNA from total DNA was performed by enzymatic digestion of linear DNA using the plasmid-safe DNase (PS-DNase) (Lucigen^®^, Epicentre, UK) that “digests linear dsDNA and, with lower efficiency, closed-circular and linear ssDNA to deoxynucleotides at slightly alkaline pH”. The circular DNA extraction protocol is described in **Figure S2**. PS-DNase digestion was carried out for 4 days with addition of fresh PS-DNase every 24 hours (4 times). This protocol was used independently with 2 sub-samples from the pooled total DNA extracted from the 5 IHHNV-positive *P. monodon* samples, but only once with a sub-sample of total DNA extracted from the 5 IHNNV-negative *P. monodon*. During the extraction process, *XhoI* enzyme in the protocol was used to cut shrimp chromosomal DNA into smaller fragments to accelerate DNA digestion by PS-DNase. The enzymes *Xho*I was chosen because it has no cutting site in shrimp mitochondrial DNA or in the IHHNV genome. If there were IHHNV-cvcDNA entities that contained portions of host DNA, they might be cut by this enzyme and be lost during circular DNA preparation. After digestion and extraction, the quantity of putative circular DNA was determined by Qubit fluorometer (Invitrogen, USA).

### 2.4 Confirmation and quality of circular DNA and circular viral copy IHHNV-DNA (IHHNV-cvcDNA)

To confirm the presence of circular DNA and lack of linear chromosomal DNA in the circular DNA preparation, PCR tests were carried out using 1) elongation factor 1 alpha (EF-1α) primers *Pm*Ef-1α-F/ *Pm*Ef-1α-R that yielded a 122 bp amplicon as a representative of chromosomal linear DNA and 2) a shrimp mitochondrial DNA (mtDNA) PCR detection method using primers *Pm*mtDNA-F/*Pm*mtDNA-R that yielded a 150 bp amplicon representing circular DNA. The presence of amplicons for 2 targets in the pre-digested DNA extract and absence of EF-1α amplicons but presence of the mtDNA amplicons in the post-digestion extract would confirm the success of circular DNA preparation. The ddPCR reaction with 309NF/309NR primers was used to quantify the IHHNV fragments in the pre-digestion DNA extract (presumed to contain non-circular IHHNV genome + IHHNV-cvcDNA) and post-digestion DNA extract (presumed to contain only IHHNV-cvcDNA).

**Figure S2.**
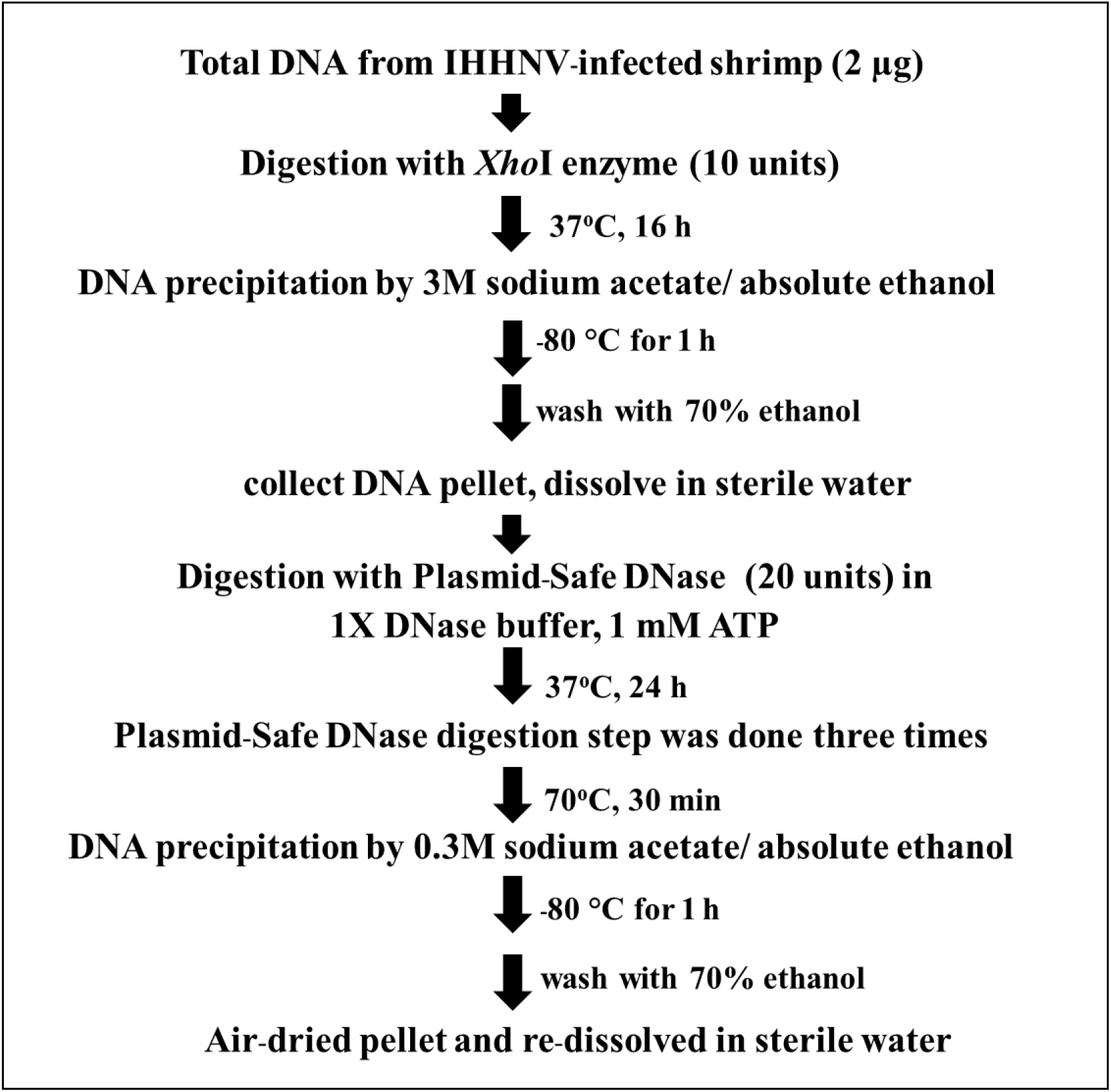
The protocol used to prepare circular DNA from the infected shrimp. Two micrograms of total DNA extract were obtained from IHHNV-infected shrimp by using commercial available DNA extraction kit (Qiagen, Germany). The total DNA was pre-digested with *Xho*I to enhance the digestion efficiency of plasmid-safe DNase in next step. This step could be replaced by an alternative *NotI* enzyme which suitable for shrimp DNA digestion. Subsequently, pre-digested DNA was extracted and further incubated with 20 U plasmid-safe DNase (PS-DNase, Lucigen^®^, Epicentre, UK) and 1mM ATP at 37 °C for 24 h. This digestion was repeated in total 4 days by addition of 20U PS-DNase and ATP in every 24h. The remaining putative circular DNA was precipitated by 0.3 M sodium acetate in absolute ethanol. The total yield of circular DNA was dissolved in DNase-RNase free water and then determined the concentration by Qubit fluorometer (Invitrogen, USA).

### 2.5 Amplification and sequencing of cvcDNA-IHHNV in the circular DNA extract

The concentration of purified circular DNA extract from IHHNV-infected shrimp was quantified by Qubit fluorometer (Invitrogen, USA). To obtain a sufficient concentration for sequencing, the circular DNA (40 ng) was subjected to rolling circle amplification (RCA) using Repli-G midi kit (Qiagen). RCA is a technique used for multiple amplification of circular DNA to obtain sufficient amplicons for next-generation nucleotide sequencing (NGS) (20).The RCA amplified products were verified by gel electrophoresis and capillary electrophoresis to determine DNA quality before sending the DNA for sequencing using the 150 bp pair-end sequencing method (Ilumina sequencing) by Novogene Co. Ltd., Hong Kong. Random fragmentation of the RCA product was by sonication. The resulting DNA fragments were end polished, A-tailed and ligated with the full-length adapters of Illumina sequencing. This was followed by further PCR amplification with P5 and indexed P7 oligos. The PCR products for the final construction of libraries were purified using the AMPure XP system. Then libraries were checked for size distribution by Agilent 2100 Bioanalyzer (Agilent Technologies, CA, USA), and quantified by real-time PCR (to meet the criteria of 3 nM). For data analysis, the raw reads of nucleotide sequences were *de novo* assembled and compared to IHHNV reference genomes in databases. The 21-nt clean reads was mapped with two IHHNV reference genome sequences, i. e., DQ228358 and AF218266 by bowtie2 and then average counts for all reads from 21-nt position along the virus reference sequences were plotted. The plot of mean reads data was constructed by ggplot in R program (https://www.r-project.org/).

### 2.6 Confirmation of the circular form by tail joining amplification

Primers were designed based on the cvcDNA sequencing result to prove that the annotated circular DNA sequences obtained were present in the original cvcDNA sample preparations. Since the sequences and contigs obtained from NGS were reported as linear nucleotide chains, it was necessary to confirm the presence of circular forms in the original circular-DNA extract. This was done by outward facing primers designed specifically from the 3’ and 5’ ends of the linear dsDNA fragments such that PCR amplification would produce amplicons only from a matching cvcDNA sequence, but not from a matching linear fragment. Specifically, either 3’ primer IHHNV3031F or IHHNV3766F was used together with a ring-closing reverse 5’ primer IHHNV128R (**Figure 2**). This is a standard protocol to test for circular dsDNA in *de novo* sequencing of dsDNA viruses (21). Single step PCR was performed using a One-*Taq*^™^ PCR reaction kit (NEB, USA) with 30 amplification cycles and 2 ng of original circular-DNA extract as the DNA template. The general protocol for PCR was 94°C for 5 min followed by PCR for 30 cycles of 94°C for 15 s, 55°C for 15 s, 72°C for 30 s and then 72°C for 5 min. The PCR products were determined by 1.5% agarose gel electrophoresis and ethidium bromide staining. The amplicon band was cut and purified from the agarose gel before cloning into pGEM-T easy vector (Promega, USA). The plasmids were transformed into *E. coli* DH5α. The plasmids prepared from 8 clones (no. 1-8) were selected and subjected to digestion with *Eco*RI enzyme to check for the variation of inserted IHHNV fragments. Four of these products from clones no. 2, 4, 6, and 7 were sent for sequencing using T7/SP6 primer (Macrogen, Korea).

**Figure 2.**
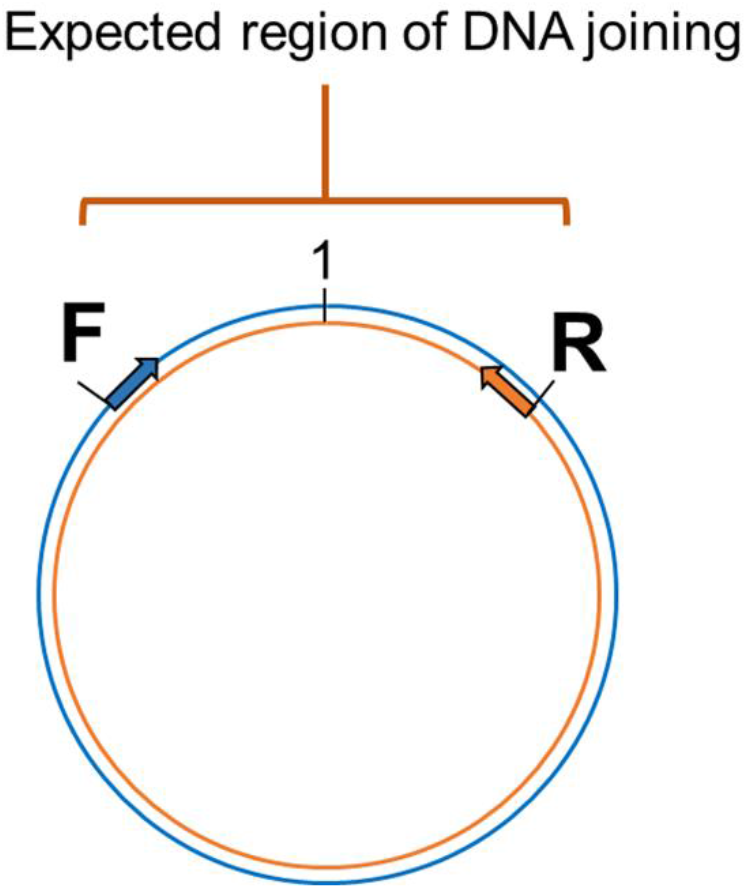
Diagram showing the method to prove circular DNA forms of IHHNV-cvcDNA by PCR amplification using outward-facing forward and reverse primers indicate the designed primers regions base on assembled linear nucleotide sequences derived from the putative cvcDNA sequencing. Single step PCR amplification was carried out using purified cvcDNA as a template. The occurrence of positive PCR amplicons indicated closure of circular DNA including the nucleotide sequences for the end-joining region.

### 2.7 Inhibition of IHHNV replication by IHHNV-cvcDNA in shrimp challenge tests

A batch of juvenile *Penaeus vannamei* (2-3 g body weight, n= 50) were obtained from the shrimp demonstration farm in Chachoengsao province and maintained in the laboratory in a 500L tank containing artificial seawater (15 ppt salinity) for 2-days with continuous aeration and water temperature between 28-30°C. The shrimp were fed with commercial feed pellets at 5% body weight daily until starting experiments. Prior to experiments, a sub-sample of 3 arbitrarily selected shrimp was tested by PCR for the absence of IHHNV using the short-amp IHHNV detection method. This would later be compared with the negative test results expected from the negative control group shrimp at the end of the IHHNV challenge experiment. The shrimp were divided into 3 groups; the PBS injection group (negative control, n= 5), the IHHNV injection group (positive control, n=5), and the test group injected with 100 ng circular DNA extract + crude IHHNV stock (n=10). The crude IHHNV stock was diluted with 1X cold PBS pH 7.4 to obtain 1 x 10^7^ copies/ 50 μl to inject individual shrimp in the IHHNV- injected positive control group and cvcDNA test group. In the test group, the diluted virus was mixed with the circular DNA preparation containing putative IHHNV-cvcDNA before intramuscular injection into individual shrimp, while the negative control group was injected with 50 μl PBS only. At day 5 post-injection, shrimp pleopods were collected from individual shrimp and then subjected to total DNA extraction followed by total DNA assessment by NanoDrop spectrophotometer (Thermo scientific, USA). Then, 100 ng was used as the template for PCR analysis for IHHNV replication level using the long-amp IHHNV method.

PCR intensities were determined using the Gel Doc^™^ EZ Gel Documentation System (Bio-Rad, USA). The relative ratio of virus level was calculated corresponding to the internal control gene expression (Ef-1α). Differences in IHHNV replication were determined by calculating the mean relative Ef-1α amplicon band intensity in the agarose gels followed by appropriate adjustment of the IHHNV band intensities before comparison of by One-Way ANOVA. Differences were considered significant at *p* ≤ 0. 05. Data analyses and graph preparations were carried out using using GraphPad Prism version 7. 0 (https://www.graphpad.com/scientific-software/prism/).

## 3. Results

### 3.1. Putative circular DNA was extracted from IHHNV-infected shrimp

After total DNA extracted from IHHNV-infected and non-infected shrimp (*P. monodon*) from the same source were exposed to PS-DNase digestion for 4 days, the EF-1α amplicon linear DNA marker could no longer be detected, in contrast to the untreated control. This indicated that all linear DNA had been digested. At the same time, the positive-control, circular-mtDNA could still be detected, indicating that circular DNA constructs could survive the exonuclease treatment (**Figure 3**).

**Figure 3.**
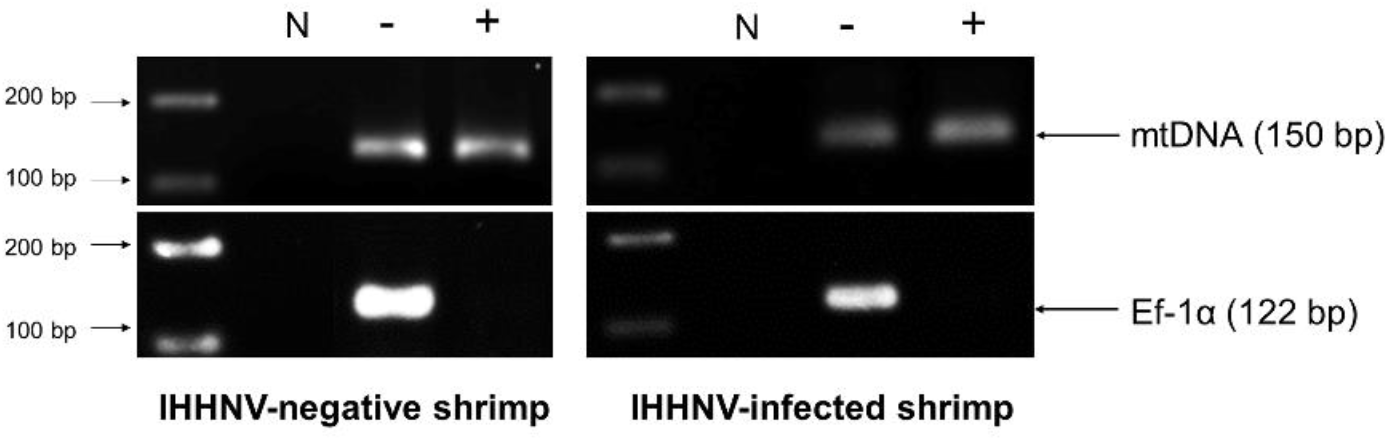
Photographs of agarose gels show amplicons from total DNA extracts of IHHNV-negative shrimp (*P. monodon*) and IHHNV-infected samples before digestion and after digestion without (-) and with both *XhoI* and PS-DNase digestion (+), respectively. PCR detection using mitochondrial DNA (mtDNA) primers and EF-1α primers as markers showed loss of an EF-1α amplicon after digestion but retained presence of the amplicon for circular mtDNA for both preparations.

Note that the band intensity for the circular-mtDNA amplicon was less than that from initial DNA preparation prior PS-DNase digestion. This may have resulted from the PCR conditions that used different concentrations of DNA template or to loss of DNA during the precipitation and recovery step or possibly to partial digestion of the circular DNA itself. However, this did not negate it as a circular-DNA marker.

Starting from 2 μg of total DNA extracted from IHHNV-infected shrimp (*P. monodon*), there remained a putative circular DNA concentration in the range of 20-40 ng in the DNA extract after 4-day digestion (a reduction of 98 to 99%). Using digital droplet PCR to measure the quantity of IHHNV in the pre-digestion and post-digestion preparations (**Figure 4**) revealed that the average pre-digestion IHHNV quantity of 3.0 x 10^5^ copies/ng DNA had dropped to 1.7 x 10^3^ copies/ng DNA. This constituted a residual of approximately 0.6 % of the initial IHHNV-DNA quantity after linear DNA digestion (i. e., 99.4% reduction). From a repeated digestion with a parallel sample of the same IHHNV-infected shrimp DNA extract, the final IHHNV quantity in the putative cvcDNA preparation was 1.2 x 10^3^ copies/ ng DNA and was not significantly different from the quantity in the first preparation (*Student’s* T-test, *p* = 0.37). IHHNV could not be detected in the negative control digests obtained from IHHNV-negative shrimp.

**Figure 4.**
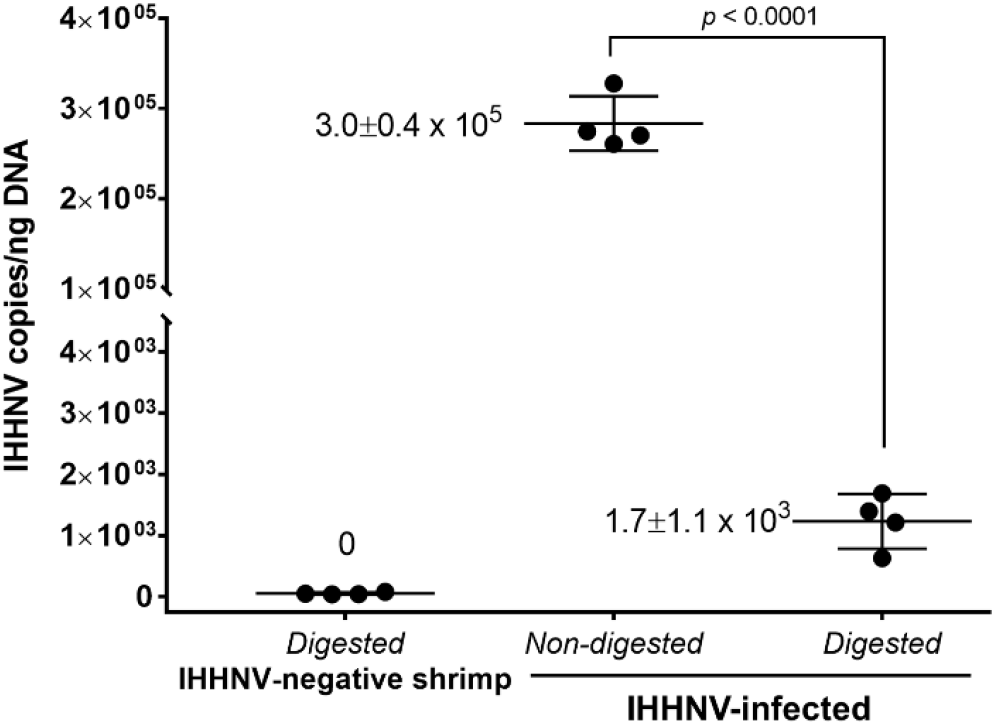
Detection of IHHNV by the digital droplet PCR method in non-digested and digested total DNA prepared from IHHNV-negative and IHHNV-infected shrimp (*P. monodon*).

In addition, exposing the putative cvcDNA extract to the restriction enzyme *Hpa*I specific for IHHNV (but not active for shrimp mitochondrial DNA) resulted in a negative PCR test for IHHNV but a retained positive PCR test result for mtDNA (**Figure 5**). This supported the contention that the IHHNV-positive PCR test result arose from IHHNV-cvcDNA. In summary, the results from Figs. 4 and 5 suggested that the IHHNV copies detected in the DNA extract from the PS-DNase digestion mix from IHHNV-infected *P. monodon* consisted of residual IHHNV in the form of cvcDNA.

**Figure 5.**
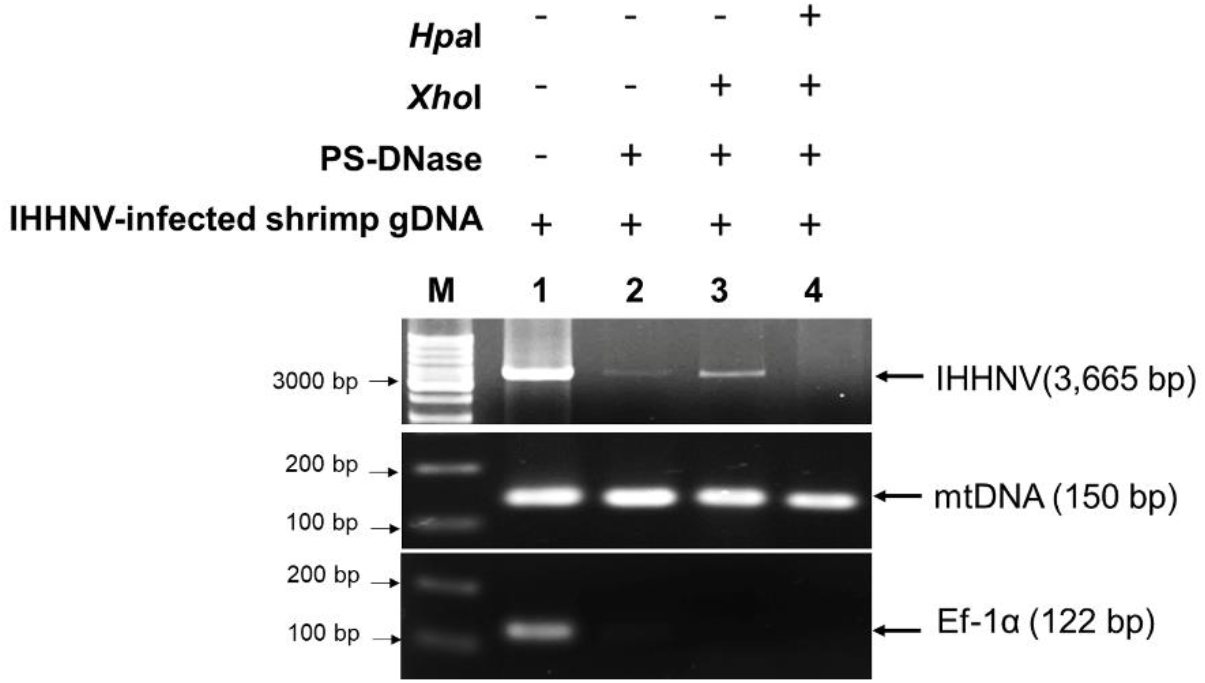
Photographs of agarose gel confirmation of IHHNV-cvcDNA presenting in putative circular DNA extract. The putative circular DNA extract in first preparation was exposed to specific IHHNV genome digestion enzyme, *Hpa*I and then the IHHNV genome sequence was determined by PCR. The result showed that absence of PCR amplicon was observed post-HpaI digestion. Whereas, mtDNA representing survive circular DNA was remained positive.

### 3.2. Putative IHHNV-cvcDNA suppressed IHHNV replication

Before investing in the cost of NGS sequencing, experiments were conducted to determine if the IHHNV putative cvcDNA preparation could protect shrimp against infectious IHHNV in a laboratory challenge test. The details of IHHNV inoculum preparation from our *P. monodon* samples and testing of its infectivity in *P. vannamei* are given in **Figure S3**.

**Figure S3.**
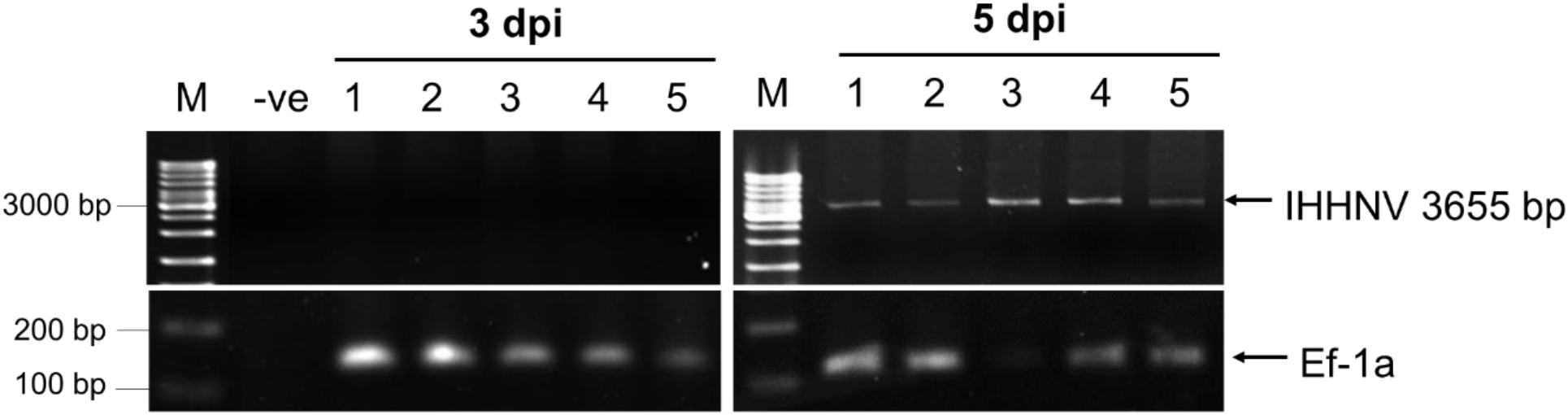
Infectivity testing of the crude IHHNV stock. To test infectivity of the IHHNV stock derived from IHHNV-infected shrimp, 10 naïve *Peneaus vannamei* were injected intramuscularly with 50 μl of the diluted stock (1 x 10^7^ copies of IHHNV). At days 3 and 5 post injection, gills from 5 shrimp were collected arbitrarily and their genomic DNA was extracted and subjected to the long-amp PCR analysis method to determine IHHNV genome replication. Five shrimp injected with the stock gave negative test results for IHHNV on day 3 but positive test results on day 5, indicating IHHNV replication and confirming the infectivity of the stock.

On day 5 post-injection in the protection test, *P. vannamei* from all 3 groups were collected and the DNA was extracted and checked by PCR using the long amp- IHHNV detection method with EF-1α as the internal control (**Figure S4**).

**Figure S4.**
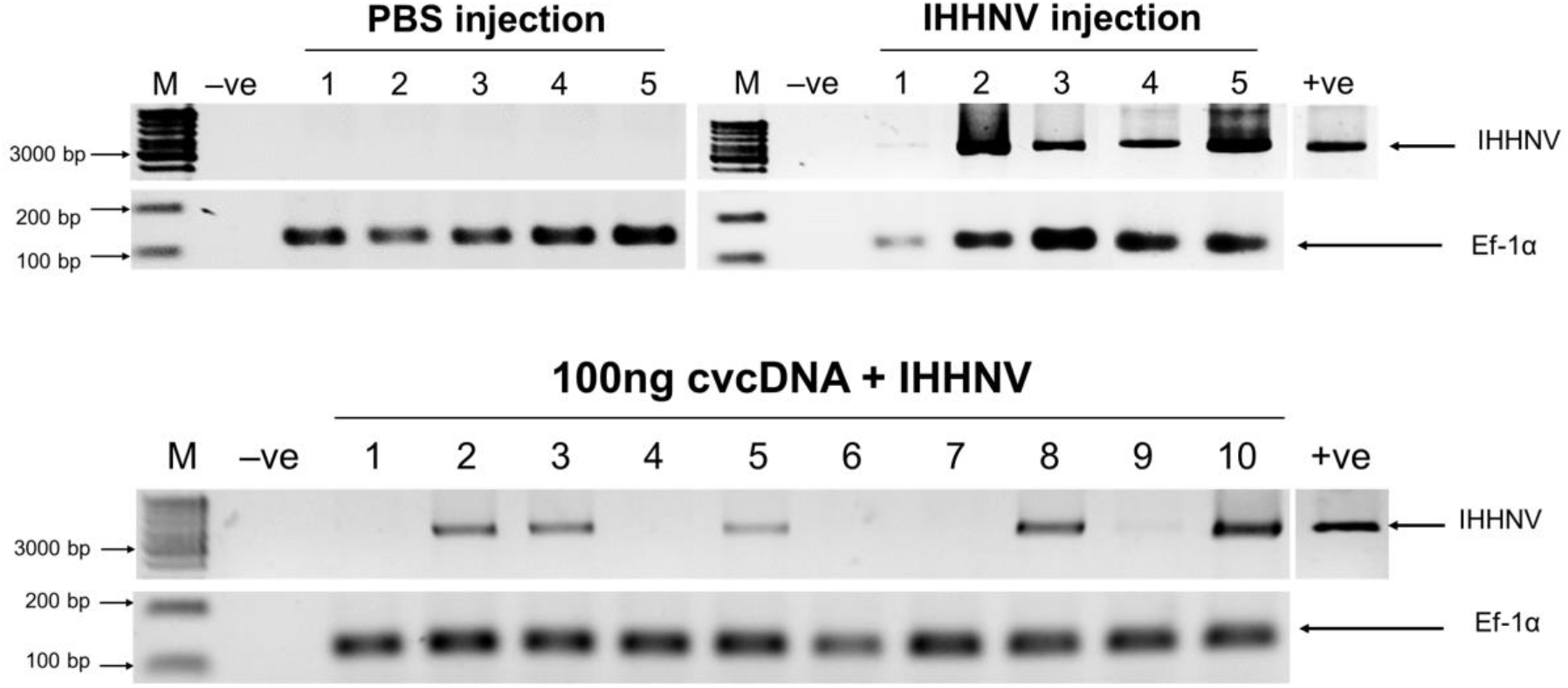
Photographs of inverted agarose electrophoresis gel showing IHHNV PCR amplicons from *P. vannamei* challenged with IHHNV. Long-amp PCR analysis indicating 3665 bp genomic DNA of IHHNV viral genome replication was seen only in shrimp injected with IHHNV (i. e., not the shrimp injected with PBS). Band intensities for the internal control Ef-1α were averaged and the average was used to adjust IHHNV band intensities before statistical comparison using one-way ANOVA.

The relative intensities of amplicon bands adjusted by the mean average of EF-1α intensities were compared (**Figure 6**). The 5 shrimp in the negative control group injected with PBS gave no PCR amplicons for IHHNV, while the positive control group of 5 shrimp injected with IHHNV gave a mean band intensity of 1.2. In contrast, the group of 10 shrimp injected simultaneously with IHHNV and 100 ng of putative IHHNV-cvcDNA gave a mean amplicon intensity of 0.2 that was significantly lower intensity (*p*<0.01) than the positive control by a One-Way ANOVA test. We considered this sufficiently encouraging to proceed with NGS sequencing.

**Figure 6.**
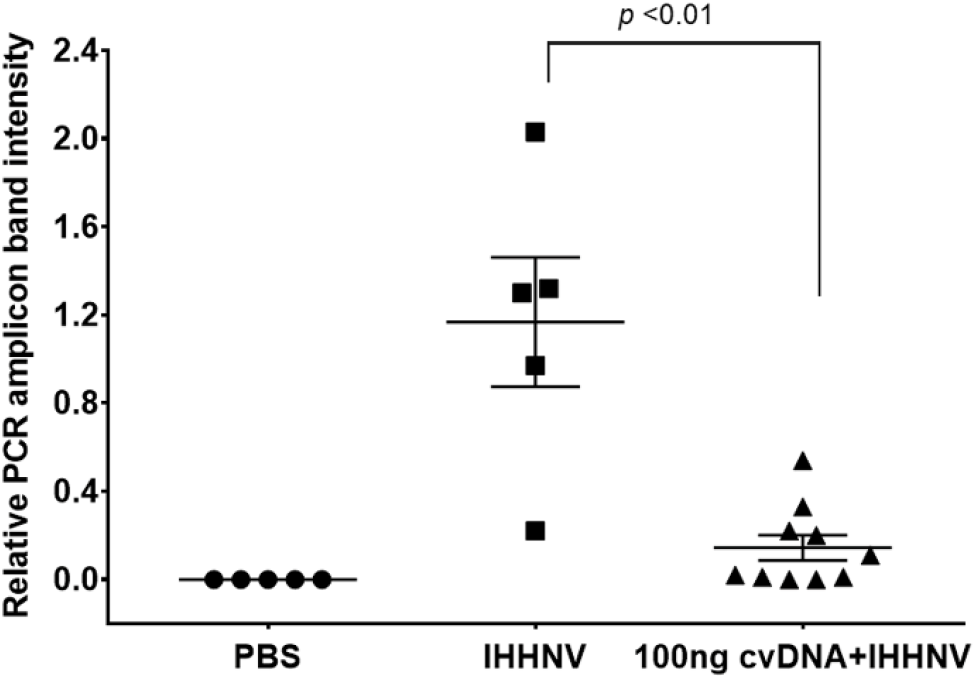
Graph of average band density for IHHNV-PCR amplicons among shrimp (*P. vannamei*) injected with IHHNV only or with IHHNV plus putative IHHNV-cvcDNA. PBS indicates IHHNV detection results for the naïve shrimp negative control.

### 3.3. RCA amplification of IHHNV-cvcDNA

The crude cvcDNA preparation (40 ng) was subjected to rolling circle amplification (RCA) using an REPLI-g mini kit (Qiagen) followed by quantification with a Qubit fluorometer revealing a total yield of 40 μg of RCA-amplified product. Subsequent agarose gel and capillary electrophoresis revealed that the majority of the RCA amplicons were of sizes larger than 11 kb (**Figure S5**). After sequencing of the RCA products, total reads of approximately 8 Mb nucleotides (8,363,236 bases) were obtained based on Illumina DNA sequencing with 99% effectiveness. A low sequencing error rate (0.03%) was found, as shown in **Table 2**. *De novo* assembly of the raw read sequences gave 81,026 contigs. A detailed analysis of the assembly is shown in **Table 3**.

**Table 2.** Sequencing information from the cvcDNA preparation obtained using the Illumina sequencing platform.

| Sample | Raw reads (bases) | Raw data (Gb) | Effective (%) | Error (%) | Q20 (%) | GC (%) |
| --- | --- | --- | --- | --- | --- | --- |
| IHHNV-cvcDNA | 8,363,236 | 1.3 | 99.73 | 0.03 | 96.79 | 47.1 |

**Table 3.** Statistics of the assembled contigs from NGS sequence analysis.

| Assembly | Total number of assembly (contigs) |
| --- | --- |
| # contigs | 81,026 |
| # contigs ($\leq 1,000$ bp) | 4,960 |
| # contigs (1001-5,000 bp) | 307 |
| # contigs (5001 to 10,000 bp) | 132 |
| GC (%) | 42.18 |
| Reference GC (%) | 42.98 |
| N50 | 553 |
| L50 | 17,163 |

**Figure S5:**
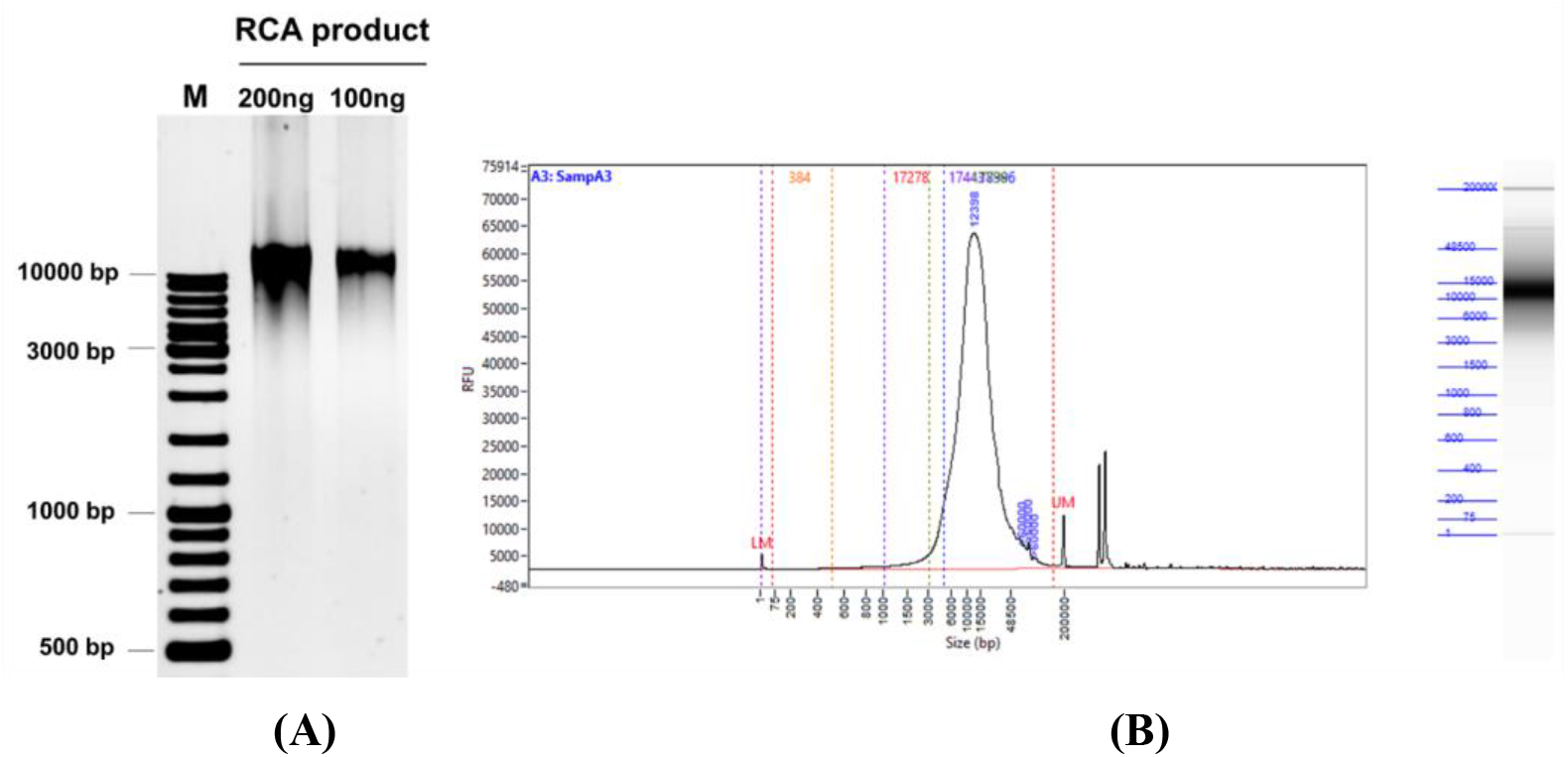
Profiles of the RCA-amplified products from the cvcDNA template preparation as determined by electrophoresis. **(A)** Photograph of a 1.5% agarose electrophoresis gel. **(B)** Graph of the output from capillary electrophoresis. Results from both techniques revealed that the majority of the products had sizes over 11,000 bp.

Diagrams of the distribution of cvcDNA sequence reads related to IHHNV reference sequences were plotted. Alignment of the frequency reads in units of 21-nt bites against GenBank records for IHHNV revealed high homology to 2 accession numbers, **i.e.** GenBank accession no. AF218266 and GenBank accession no. DQ228358 (**Fig. S6**). Short read DNA assembly based on the reference IHHNV sequences was carried out and the 3 longest contigs were selected for deeper analysis. Two of these (NODE_444 and NODE_1) matched the IHHNV genome accession no. AF218266 (**Fig. S7A**) from an extant, infectious type of IHHNV. The third contig (NODE_439_3766) matched accession no. DQ228358 which contains an ancient IHHNV sequence (**Fig. S7B**) that is inserted into the genome of some *P. monodon* specimens. Thus, it should now be called an endogenous viral element or EVE. It was discovered before use of the term EVE and was called “ non-infectious” IHHNV (19). This was a completely unexpected and unpredicted result and was thus, not included in our experimental hypothesis for this study.

**Figure S6.**
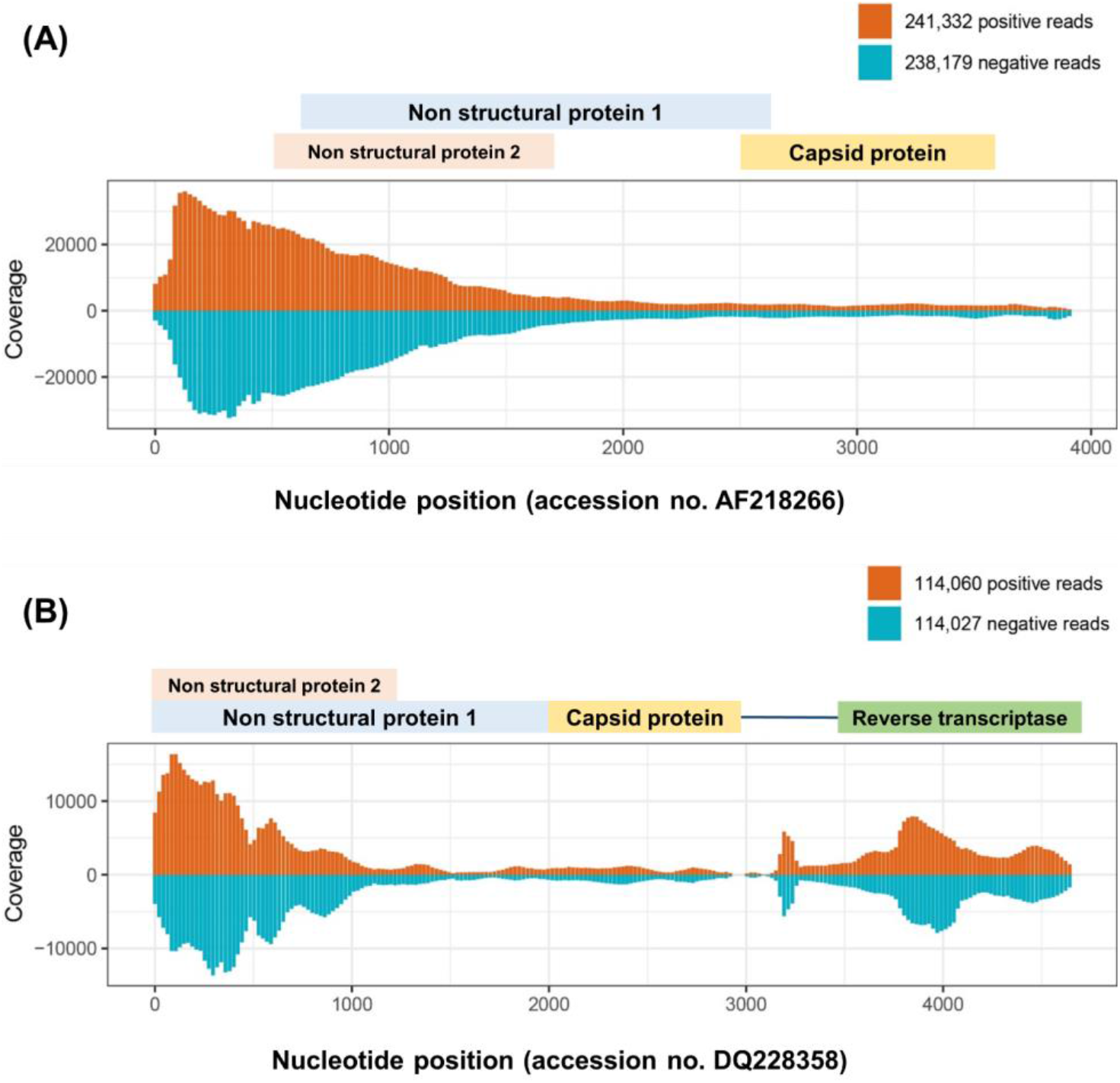
Diagrams of cvcDNA sequence reads distribution related to IHHNV reference sequences. The bar plots indicate distribution of the mean count of sequence reads obtained from DNA sequencing. The 21-nt sequence reads are shown as both plus and minus reads throughout the genome length. Distribution of 21-nt reads related to **(A)** GenBank accession no. AF218266 and **(B)** GenBank accession no. DQ228358. The shaded boxes above the graph represents the open reading frames of the related sequences corresponding to their nucleotide positions.

**Figure S7.**
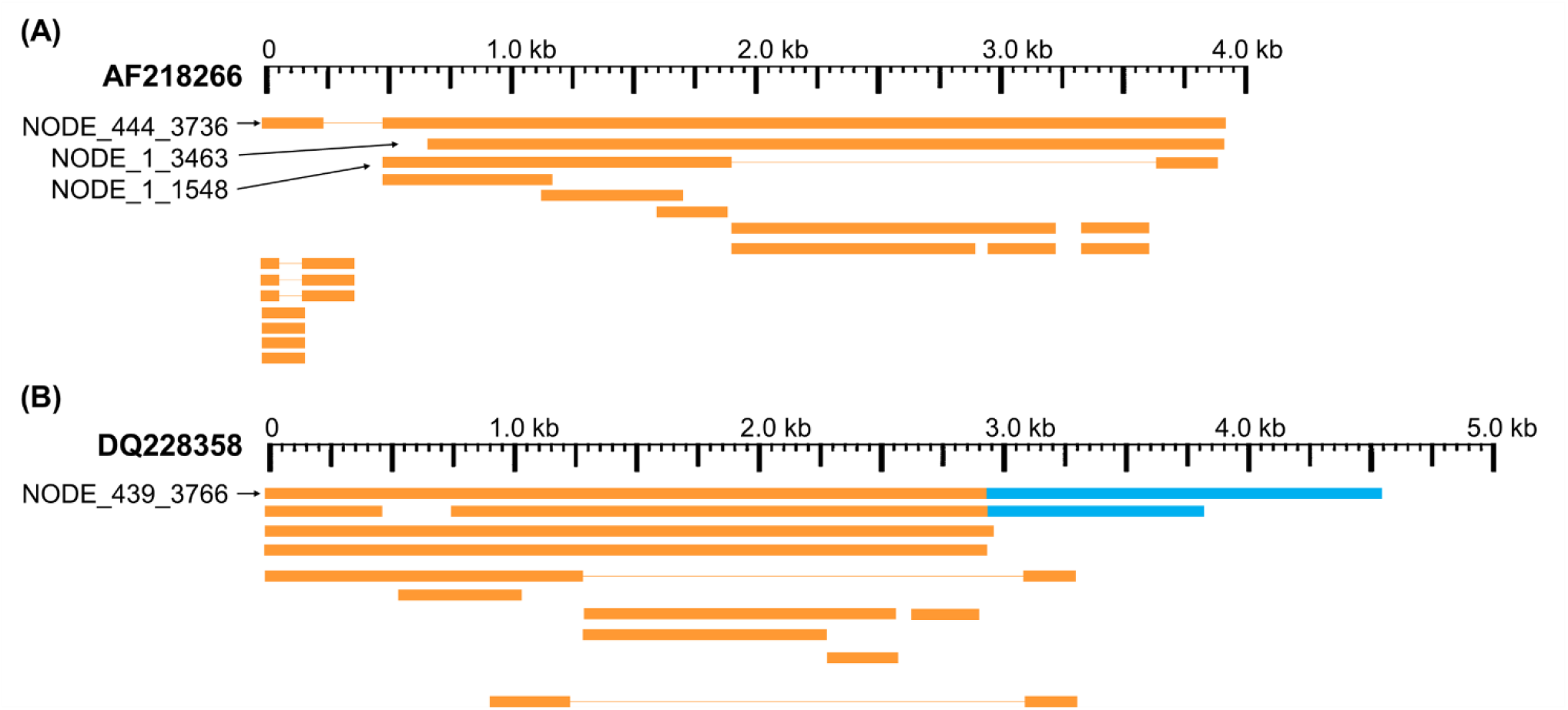
Schematic diagram showing the sequence similarity compared between putative cvcDNA contig sequences and Genbank records. The scale bar indicates nucleotide positions. Each box represents the sequence of an individual DNA contig. In the lowest cvcDNA in **A**, there is a long deletion (indicated by a line) when compared to the GenBank record. **(A)** cvcDNA sequences with high similarity to the IHHNV virus GenBank record AF218266. **(B)** Sequences with high similarity to non-infectious IHHNV GenBank record DQ228358. The regions in blue in **B** indicate the portion of the DQ228358 sequence that is part of a host shrimp transposable element and relates to the insertion point of the EVE in the shrimp genome.

NODE_444 and NODE_1 showed 98 and 99% identity, respectively, to AF218266 (**Table 4**) and 99% sequence identity to the IHHNV in the frozen *P. monodon* we used for preparation of the putative cvcDNA extract. These results were consistent with our prediction that IHHNV-cvcDNA would arise after IHHNV challenge. NODE_439_3766 showed 98% sequence identity to DQ228358.

**Table 4.** List of longest contigs with similarity to IHHNV genome references.

| Sequence ID | Length (bp) | Backbone similarity |
| --- | --- | --- |
| NODE_439_3766 | 3,766 bp | DQ228358 (98%) |
| NODE_444_3736 | 3,736 bp | AF218266 (99%) |
| NODE_1_3463 | 3,463 bp | AF218266 (99%) |
| NODE_1_1548 | 1,548 bp | AF218266 (99%) |

### 3.4. PCR confirmed that contigs matching AF218266 arose from cvcDNA

Outward facing primers 3031F/ 128R designed to match the ends of the linear contigs NODE_1_3463 and Node_444_3736 obtained by NGS (**Figure 7A**) gave rise to PCR amplicons, confirming that the linear contigs obtained by NGS sequencing from **Table 4** arose from cvcDNA. Failure to obtain amplicons would have indicated that the target sequence was a linear DNA fragment. The PCR results gave a single PCR amplicon of approximately 1,500 bp (**Figure 7B**). This confirmed that the contigs ID NODE_1-3463 and/ or NODE_444-3736 were derived from closed-circular DNA forms. The band of the 1,500 bp amplicon was purified and cloned. The plasmids from each clone were digested with *Eco*RI enzyme and the digestion results are shown in **Figure 7C.** There were variations in the amplicon sizes among the 8 selected clones indicating a mixture of amplicons. Four selected clones were subjected to plasmid sequencing. Approximately 600-800 bp were read using forward and reverse primers and the assembled sequences (approximately 1,000-1,500 bp) matched sequences in the IHHNV reference genome AF218266. They corresponded to the expected amplified regions based on the GenBank reference genome. The ring in **Figure 7D** represents a model cvcDNA of variable overall length with the orange portion indicating the contigs for NODE_1_3463 or NODE 444-3736 (or similar contigs that might contain the targets for primers IHHNV3031F/IHHNV128R at each end). In other words, the amplified sequences linked the two ends of each contig to form a circle and matched the expected corresponding sequences in the reference genome (**Figure 7E**).

**Figure 7.**
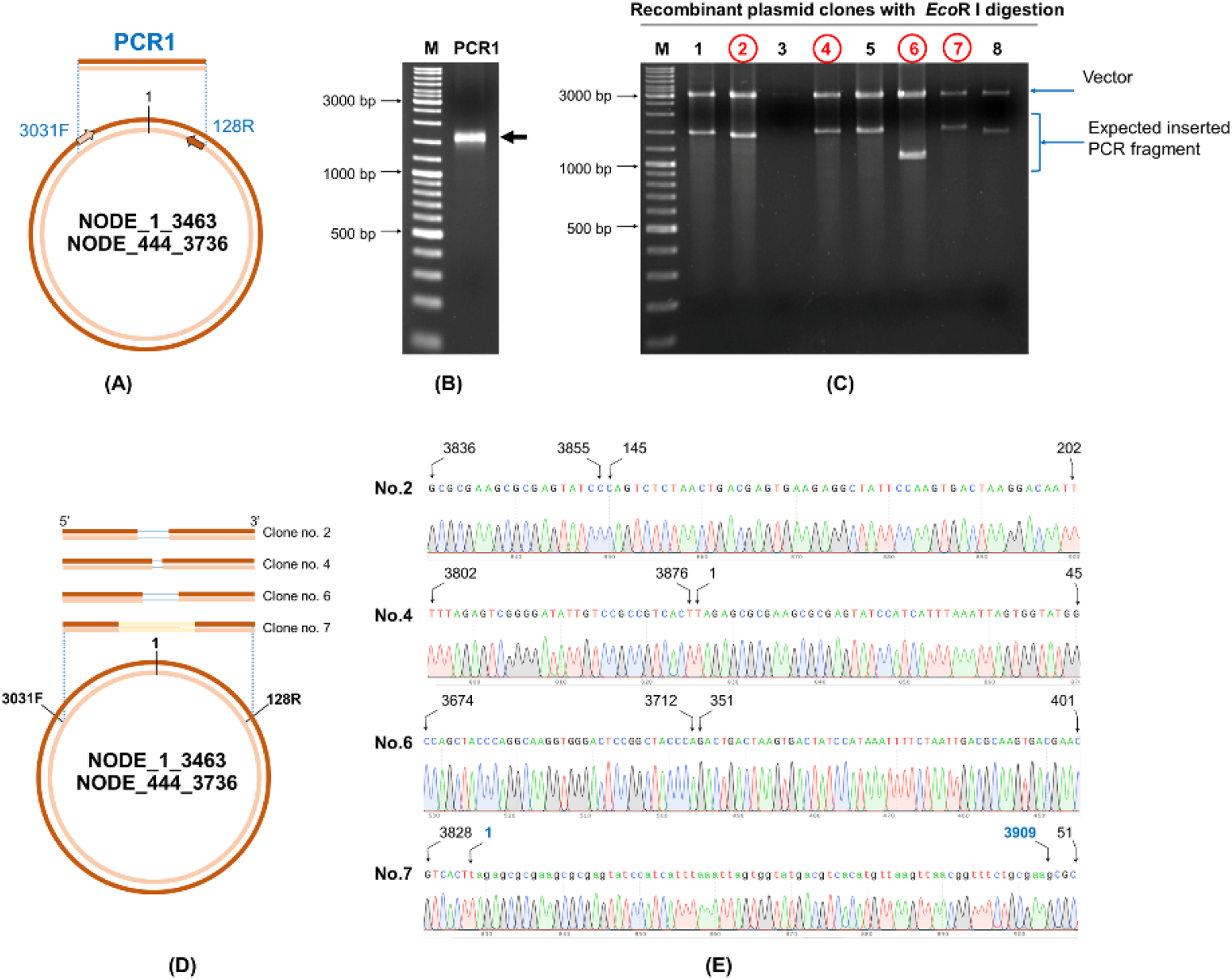
Confirmation of IHHNV-cvcDNA sequences homologous to GenBank record AF218266 by PCR and DNA sequencing. **(A)** A diagram using ID NODE_1-3463 as a model with primer positions (IHHNV3032F/IHHNV128R) and amplification directions indicated. **(B)** Photograph of an electrophoresis gel showing the amplified broad band of approximately 1,500 bp that was obtained from use of primers IHHNV3031F/IHHNV128R. **(C)** *Eco*RI digestion of the plasmid preparations from 8 selected clones. The plasmid of clone no. 2, 4, 6, and 7 (in red) were subjected to sequencing using T7/ SP6 primers. **(D)** Diagram showing PCR sequence reads and alignments of the cvcDNA variants corresponding to the reference contigs no. NODE_1_3463 or NODE 444-3736. The ring represents a model cvcDNA of variable overall length with the orange portion indicating the contigs for NODE_1_3463 or NODE 444-3736. The orange boxes above the circle indicate variations in contiguous, amplified closure lengths that also share 99% identity to AF218266. The lines within some of the boxes simply indicate which missing part of the match to AF218266 has resulted in the smaller closure length. **(E)** Junction analysis based on PCR fragment sequencing in **7C** showing a chromatogram for continuous DNA reads. None of the closed circles contained a match to a whole IHHNV genome sequence. Numbers indicate nucleotide position corresponding to reference genome accession no. AF218266.

### 3.5. PCR confirmed that contigs matching DQ228358 arose from cvcDNA

Next we confirmed that contig no. NODE_439_3766 that matched the GenBank record DQ228358 also arose from cvcDNA. This contig must have arisen from an EVE in the genome of the experimental shrimp used, and it contained a portion of the host shrimp retrotransposon sequence to which the EVE is linked in the shrimp genome. To close the cvcDNA circle, we followed the same protocol that was followed in the section 3.4 for ID NODE_444-3736 and NODE_1-3463.

Primers IHHNV3766F/IHHNV128R designed to match the ends of NODE_439_3766 were used (**Figure 8A**). PCR results revealed an expected amplicon size of approximately 1,200 bp (**Figure 8B**). After PCR cloning, 8 positive clones showed a variety of amplicon sizes indicating several cvcDNA types (**Figure 8C**), similar to the phenomenon previously observed for ID NODE_444-3736 and ID NODE_1-3463, described in section 3.4 above. DNA sequence reads of 6 clones were of 2 general types, one showing sequence reads containing only IHHNV portions of the DQ228358 sequence (clone no. 1, data not shown) and others (clones no. 2,7) containing the DQ228358 sequence joined to its retrotransposon part (**Figure 8D-E**). Again, based on the publications linked to reference sequence accession no. DQ228358, these cvcDNA must have originated from the IHHNV-EVE called DQ228358 and its linked host-transposon sequence. This was an unexpected finding providing evidence that cvcDNA can be generated from not only cognate viruses but also from EVE in the host genome itself.

**Figure 8.**
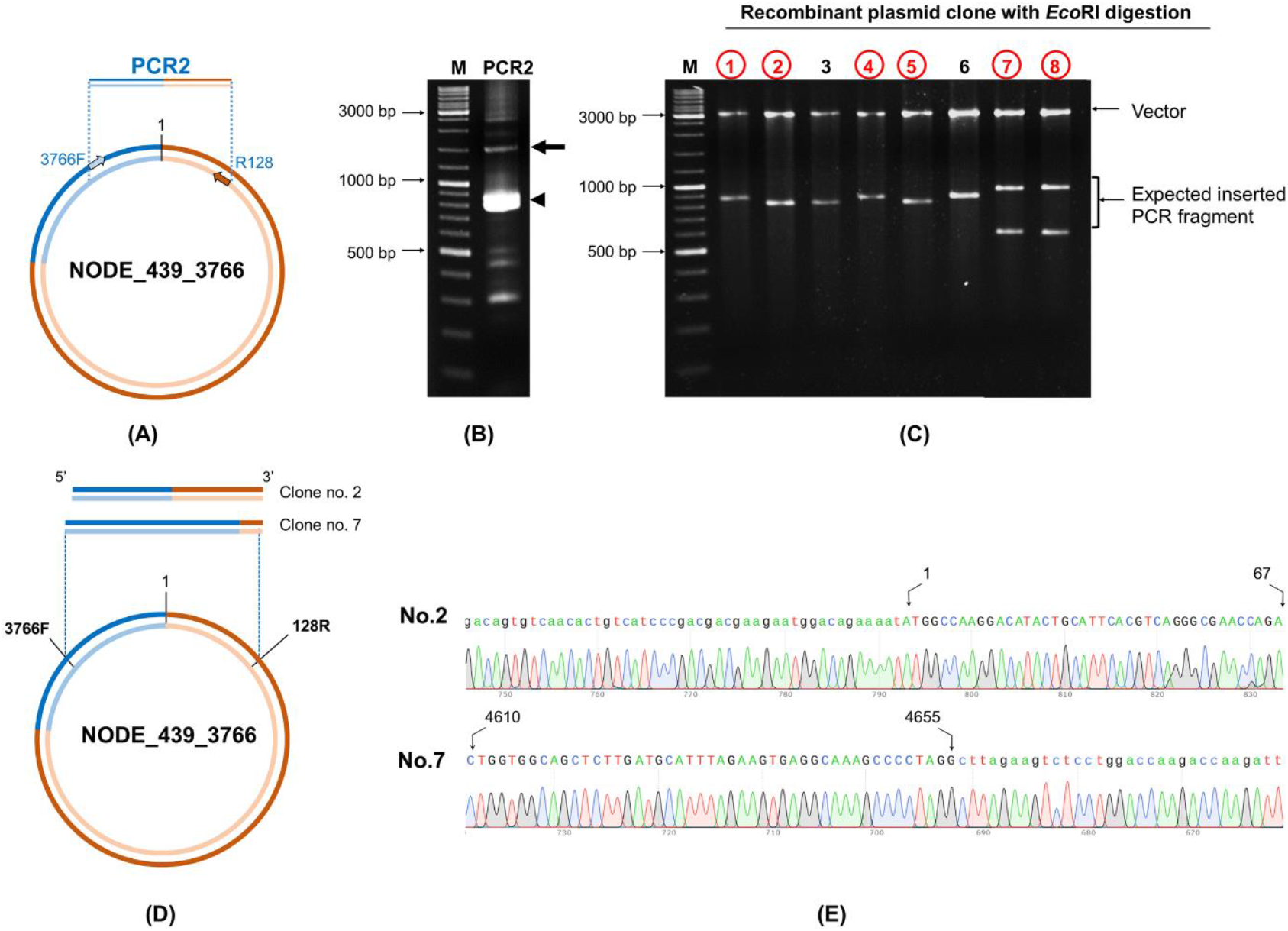
Confirmation of cvcDNA sequences of NODE_439_3766 homologous to GenBank record DQ228358 by PCR and DNA sequencing. **(A)** Diagram showing the primer pairs designed for cvcDNA amplification. **(B)** The PCR amplicon was obtained from shrimp cvcDNA using single step PCR and confirming that the annotated cvcDNA sequences were closed circular forms. The expected PCR amplicon of approximately 1200 bp was observed (arrow). However, the majority of PCR amplicons were of smaller sizes (indicated by an arrowhead). **(C)** PCR fragment ligation and cloning to a plasmid vector revealed variation in the inserted PCR fragments. **(D)** Diagram showing the PCR sequence reads and alignments of the cvcDNA variants corresponding to the reference contigs no. NODE_439_3766. **(E)** Junction analysis of DNA sequencing obtained from PCR fragment sequences in 8C showing continuous DNA reads in the chromatogram linking the 3’ and 5’ ends of the reference sequence accession no. DQ228358. Extension of the shrimp retrotransposon was also observed (additional sequences indicated by lowercase letters).

Since we made pooled DNA extracts and cvcDNA preparations from one group of IHHNV-positive and one group of IHHNV-negative specimens (*P. monodon*) obtained from the same batch of frozen shrimp, it was of interest to determine whether the IHHNV-negative preparations were also positive for the presence of EVE-DQ228358. Subsequent PCR testing revealed that the DNA extract and cvcDNA preparation derived from the IHHNV-negative shrimp (*P. monodon*) gave positive PCR test results for the DQ228358 target sequence (data not shown). We did not pursue this further by NGS sequencing, but it suggested that the IHHNV-negative shrimp (*P. monodon*) carried EVE-DQ228358 and that IHHNV infection was not necessary for it to produce cvcDNA from it.

## 4. DISCUSSION

In this study, the procedures previously published for RNA viruses in insects (Tassetto et al., 2017) were followed to prepare cvcDNA from shrimp infected with the DNA virus IHHNV. Although the detailed mechanism by which cvcDNA arises in insects was not elucidated, it was suggested to arise from the activity of host RT on RNA from the infecting RNA virus. In the case of the DNA virus IHHNV, we hypothesized that mRNA from IHHNV would be the substrate for host RT to produce cvcDNA. It might be argued that the procedures we used to prepare cvcDNA did not work and that the positive PCR results we obtained for IHHNV after following the published cvcDNA isolation protocol arose not from cvcDNA but from residual IHHNV ssDNA or from IHHNV “ rolling hairpin” intermediate replication forms (RF) (22). However, according to the manufacturer, Lucent^®^ Plasmid-Safe DNase (PS-DNase) “digests linear dsDNA to deoxynucleotides at slightly alkaline pH and, with lower efficiency, closed-circular and linear ssDNA”. Thus, the 4-day digestion should have removed IHHNV ssDNA and RF intermediates. This contention was supported by the 99.4% reduction in IHHNV copies after the digestion process. It was also supported by the loss of IHHNV detection but not mtDNA detection by PCR after the cvcDNA preparation was treated with an IHHNV-specific restriction enzyme followed by further PS-DNase treatment. In addition, the presence of cvcDNA that matched EVE-DQ228358 should not have arisen from “ rolling hairpin” intermediate replication RF, because it is a host chromosomal element, not an infectious parvovirus, and as far as we know, does not replicate, but might produce RNA transcripts. According to our cvcDNA sequencing and PCR analysis, the cvcDNA forms produced in our IHHNV-infected shrimp could be classified into two major types (**Figure 9**).

**Figure 9.**
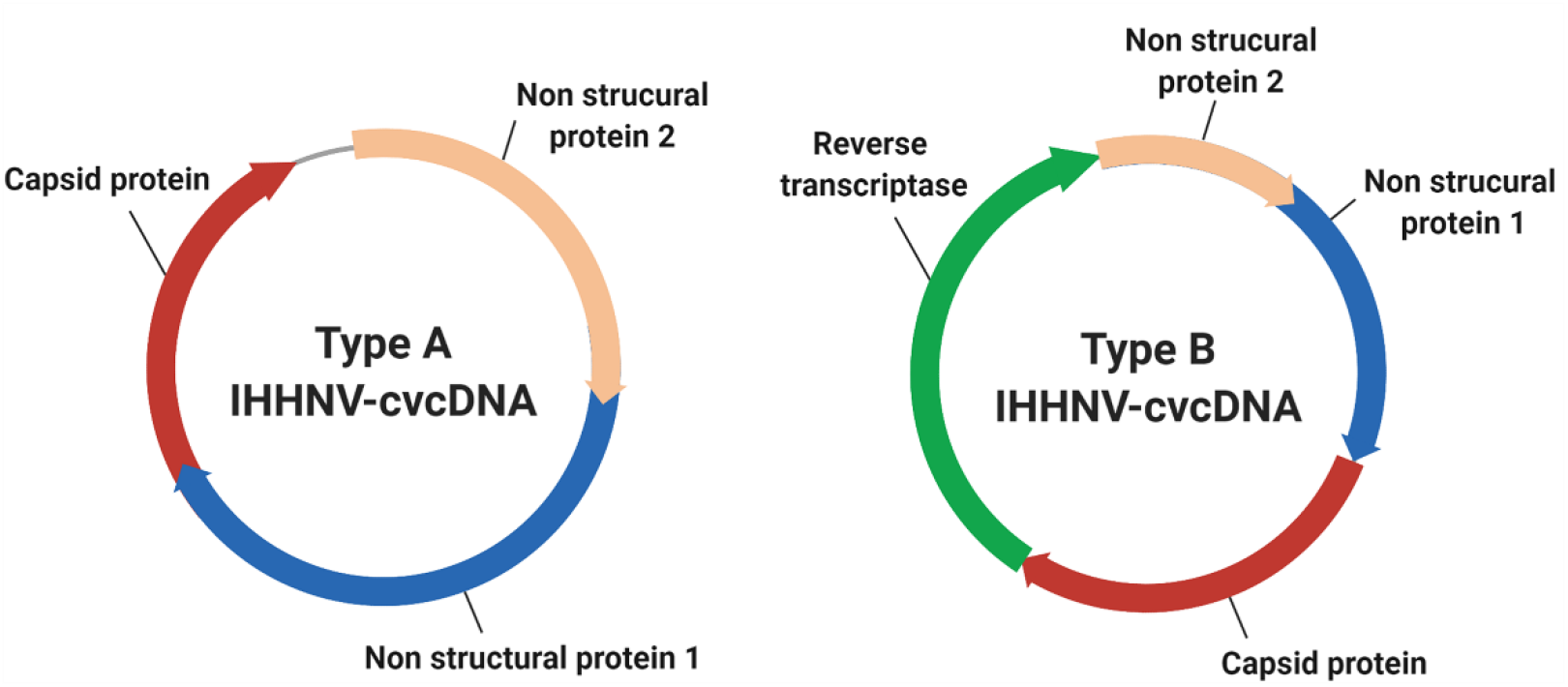
Diagram representing the two general types of cvcDNA produced in our IHHNV-infected *P. monodon*. Type A contained only DNA with sequences with 98-99% sequence identity to a currently extant species of IHHNV. Type B contained DNA with 98-99% sequence identity to an IHHNV-EVE and sometimes including (as illustrated here) a portion of the linked host DNA sequence that revealed the location of the EVE in the host genome.

The first type contained only variable-length fragments of infectious IHHNV without containing any host nucleotide sequence. All had high identity to the infectious IHHNV sequence record AF218266. We do not currently know the process by which these cvcDNA constructs were generated. However, we hypothesize that they arose from the mRNA of infectious IHHNV as a result of endogenous (host) reverse transcriptase activity, as proposed by the viral accommodation hypothesis (5) and as shown from research with RNA viruses in insects (10) (see **Figure 1**).

The second type of cvcDNA we obtained contained sequences of the EVE represented by GenBank record DQ228358. This type consisted of two sub-types, one containing only viral sequence and another containing viral sequence together with a portion of the adjacent transposable element to which it is linked in the shrimp genome. DQ228358 is known to be an integral part of the genome in some specimens of *P. monodon* from the Indo-Pacific region (19), so it was a surprising discovery, revealing that the cvcDNA may have arisen directly from the host shrimp genome via some kind of DNA polymerase. Alternatively, the EVE may have produced an RNA transcript that was subsequently used as a template by host RT that normally produces cvcDNA entities in response to viral infection (see. **Figure 1**). Given the information currently in hand, we know that EVE in shrimp (5, 14) and in insects (11) do produce RNA transcripts. We also know from the insect work that the RNA from infecting viruses can give rise to host generated cvcDNA via host RT activity. Thus, we hypothesize that EVE-DQ228358 can produce RNA transcripts that are processed in a manner similar to that used by insects, in which viral RNA serves as a template for production of lvcDNA and cvcDNA. An easy way to test this hypothesis would be to block host RT activity. The hypothesis predicts that doing so would prevent cvcDNA production from DQ228358. This pathway may be additional to the cytoplasmic processing of long, piwi-interacting-like (piRNA-like) RNA transcripts into small RNA fragments leading to RNAi via interaction with specific PIWI binding proteins, as has been shown for insects (23, 24). Whatever the mechanisms for production of cvcDNA from EVE, the extraction of cvcDNA provides a very convenient method to identify EVE in normal, uninfected shrimp and screen them for possible antiviral protective activity.

It has been previously proven that shrimp and insects can accommodate both RNA and DNA viruses in tolerated, persistent infections and that they may also carry EVE for those tolerated viruses (5). The occurrence of a protective EVE in insects has been recently proven for mosquitoes (12). Now, we have shown that cvcDNA arises in shrimp from both invading

IHHNV and from an ancient IHHNV-EVE and that the extracted cvcDNA mix could interfere with IHHNV replication. These results support the hypothesis that shrimp may have underlying mechanisms for viral accommodation for a DNA virus (1, 5) similar to those that have already been described for RNA viruses in insects.

The discovery of cvcDNA arising from EVE was unexpected and is exciting, because it means that the process of isolation and characterization of cvcDNA appears to be a convenient way to screen for the presence of potentially protective EVE in shrimp and insects. Once the cvcDNA types have been characterized, primers can be designed to identify their presence in individual specimens of breeding stocks, allowing them to be tested for protective capability. They may also be produced *in vitro* and tested for protective capability by shrimp injection in laboratory trials. EVE mimics providing the most effective protection could be amplified *in vitro* by PCR and tested in shrimp as vaccines added to feeds. Similar approaches might be used for commercial insects (e.g., silkworms and honeybees). Such protective cvcDNA could also be specifically designed and injected into the ovaries of SPF broodstock shrimp for potential insertion in the shrimp genome. Then the offspring could be screened for possession of heritable, protective EVE. Such a model suggests that it might eventually allow for the improvement of current SPF shrimp stocks to increase their range of high tolerance to serious viral pathogens. The biggest advantage is that sourcing natural and protective EVE from shrimp and insects and using them for disease control should not elicit regulatory restrictions because the vaccines and reagents used would be from natural sources and because they are non-replicative and carry no antibiotic resistance genes.

We would like to proceed with this work by producing some of the IHHNV-cvcDNA entities we have discovered *in vitro* to test for their efficacy in controlling IHHNV infections in *P. vannamei* first by injection and then (if effective by injection) by addition to feed. One major question will be whether the cvcDNA with IHHNV sequences only or those also containing transposable element sequences will be more protective. If these trials prove to be successful, a practical vaccine might be produced for IHHNV and the model developed could also be used for other viruses such as white spot syndrome virus (WSSV).

## 6. Summary

A protocol for cvcDNA preparation was used and shown to be successful for extracting IHHNV-cvcDNA that matched the sequence of infective IHHNV in *P. monodon*. The extracted IHHNV-cvcDNA was shown to inhibit IHHNV replication when it was injected into *P. vannamei* challenged with IHHNV. Subsequent next generation sequencing (NGS) of the IHHNV-cvcDNA extract revealed a variety of IHHNV-cvcDNA types, one type that originated from the infecting IHHNV and another that originated from a host EVE. This unexpected discovery of cvcDNA arising from an EVE opens the way for relatively easy identification of natural and potentially protective EVE in shrimp via cvcDNA. This may lead to applications of EVE in shrimp and perhaps insects. The detailed mechanisms related to the production of cvcDNA from infecting viruses and from EVE in shrimp remain to be revealed, but its existence constitutes a new frontier for the discovery and potential application of cvcDNA for shrimp vaccination and for improvement of viral tolerance in shrimp breeding stocks.

We declare that our discovery of cvcDNA originating from EVE constitutes a revelation of a natural process that occurs in shrimp. As such, the process of using cvcDNA to detect and study EVE cannot be considered intellectual property eligible for patenting. Thus, anyone can use this knowledge freely to screen for protective EVE via the cvcDNA they may give rise to. It is possible that during this process some specific and highly protective, natural EVE may be discovered and used directly as vaccines or regents to genetically modify SPF shrimp, but again, such discoveries and applications would not be patentable because of the natural occurrence of the EVE and the cvcDNA they give rise to. It would be tantamount to trying to patent the shrimp themselves. On the other hand, it is possible, for example, that specific inventions of non-obvious vaccines and delivery methods may be suitable for patenting.

## List of abbreviations

vcDNA: viral copy DNA(s)
lvcDNA: linear viral copy DNA
cvcDNA: circular viral copy DNA
IHHNV: Infectious hypodermal and hematopoietic necrosis disease virus
PS-DNase: Plasmid-safe DNase
EVE: Endogenous viral element(s)
siRNA: small interfering RNA(s)

## Acknowledgements

The authors would like to thank Guangdong Haid Group Co. Ltd and the National Centre for Genetic Engineering and Biotechnology (BIOTEC) of the Thai National Science and Technology Development Agency (NSTDA) for the joint funding provided to carry out this work. We would also like to thank to Dr. Donghuo Jiang for encouraging innovation, for his helpful discussions and for his efforts that enabled us to explore this frontier area of shrimp immunology.

## Ethics statement

This work followed Thailand’s laws for ethical animal care under the Animal for Scientific 83 Purposes ACT, B.E. 2558 (A.D. 2015) under project approval number BT-Animal document no. BT37/ 2563) from the National Center For Genetic Engineering and Biotechnology (BIOTEC), National Science and Technology Development Agency (NSTDA), Thailand.

## Authors contributions

Conceptualization, Data validation, Writing: ST, TWF; Methodology, Investigation, Validation: ST, PB, SS, PW, KS; Data validation, Data curation, Reviewing and Editing: ST, KS, TWF

